# Genome-wide Enhancer Maps Differ Significantly in Genomic Distribution, Evolution, and Function

**DOI:** 10.1101/176610

**Authors:** Mary Lauren Benton, Sai Charan Talipineni, Dennis Kostka, John A. Capra

## Abstract

Non-coding gene regulatory enhancers are essential to transcription in mammalian cells. As a result, a large variety of experimental and computational strategies have been developed to identify *cis*-regulatory enhancer sequences. In practice, most studies consider enhancers identified by only a single method, and the concordance of enhancers identified by different methods has not been comprehensively evaluated. Here, we assess the similarities of enhancer sets identified by ten representative strategies in four biological contexts and evaluate the robustness of downstream conclusions to the choice of identification strategy. All pairs of enhancer sets we evaluated overlap significantly more than expected by chance; however, we also found significant dissimilarity between enhancer sets in their genomic characteristics, evolutionary conservation, and association with functional loci within each context. We find most regions identified as enhancers are supported by only one method. The disagreement is sufficient to influence interpretation of GWAS SNPs and eQTL, and to lead to disparate conclusions about enhancer biology and disease mechanisms. We also find only limited evidence that regions identified by multiple enhancer identification methods are better candidates than those identified by a single method. Our results highlight the inherent complexity of enhancer biology and argue that current approaches have yet to adequately account for enhancer diversity. As a result, we cannot recommend the use of any single enhancer identification strategy in isolation. To facilitate assessment of enhancer diversity on studies’ conclusions, we developed creDB, a database of enhancer annotations designed to integrate into bioinformatics workflows. While our findings highlight a major challenge to mapping the genetic architecture of complex disease and interpreting regulatory variants found in patient genomes, a systematic understanding of similarities and differences in enhancer identification methodology will ultimately enable robust inferences about gene regulatory sequences.

## INTRODUCTION

Enhancers are traditionally defined as genomic sequences that regulate the transcription of one or more genes, regardless of orientation or relative distance to the target promoter^1^. These cis-regulatory regions can bind specific transcription factors and cofactors to increase transcription, and in current models of enhancer function, they physically interact with their long-range targets via loops in the three-dimensional chromatin structure^1–3^. Enhancers play a vital role in the regulation of genes during development and cell differentiation^3,4^. Genetic variation in enhancers has been implicated in etiology of complex disease^5,6^ and in differences between closely related species^7–9^.

Given their importance, enhancers have seen considerable study in recent years. More than 2,300 papers have been published on enhancer biology (MeSH: *Enhancer Elements, Genetic)* since the start of 2015, and hundreds of these have focused on the role of enhancers in disease. Despite the importance of enhancers, they remain difficult to identify^1,10,11^. Experimental assays that directly confirm enhancer activity are time-consuming, expensive, and not always conclusive^1,12^. And, despite recent promising developments in massively parallel reporter assays, current methods are unable to definitively identify and test enhancers on an unbiased genome-wide scale^13^. As a result, many studies have used more easily measurable attributes associated with enhancer activity, including DNA sequence motifs, evolutionary conservation, and biochemical properties, as proxies for enhancer activity^1,14–17^. For example, active enhancers localize in regions of open chromatin, which are commonly assayed by testing the sensitivity of DNA segments to DNase I nuclease followed by sequencing (DNase-seq)^18,19^. Enhancers also often have characteristic sets of histone modifications on surrounding nucleosomes. For example, monomethylation of lysine 4 on histone H3 (H3K4me1) and lack of trimethylation of lysine 4 on histone H3 (H3K4me3) is used to distinguish enhancers from promoters, while acetylation of lysine 27 on histone H3 (H3K27ac) often denotes active enhancers^1,14,20^. Additionally, features such as evolutionary sequence conservation^21–23^ and the presence of known transcription factor binding motifs^16^ or known enhancer associated proteins, such as the histone acetyltransferase p300^14,24,25^, have been used to successfully locate enhancer elements^1^. Furthermore some enhancers are transcribed, and it has become possible to map active enhancers by identifying characteristic bi-directionally transcribed enhancer RNAs (eRNAs)^26,27^, although the specificity of this pattern to enhancers has recently been questioned^28^. However, while informative, none of these attributes are comprehensive, exclusive to enhancers, or completely reliable indicators of enhancer activity. Enhancer activity is also context- and stimulus-dependent, creating an additional layer of complexity^20,29^. Furthermore, genetic variation between individuals affects epigenetic modifications and enhancer activity; however, recent studies suggest that only a small fraction (1–15%) of epigenetic modifications are influenced by nearby genetic variants^30^.

Many complementary computational enhancer identification methods that integrate these data in both supervised and unsupervised machine learning approaches have been developed^11,15,31^. Since there are no genome-wide gold standard enhancer sets, such enhancer identification studies and algorithms validate their results via a combination of small-scale transgenic reporter gene assays and enrichment for other functional attributes, such as trait-associated genetic variants, evolutionary conservation, or proximity to relevant genes. The resulting genome-wide enhancer maps are commonly used in many different applications, including studies of the gene regulatory architecture of different tissues, and the interpretation of variants identified in genome-wide association studies (GWAS)^5,6,32^. However, in these applications, a single assay or computationally predicted enhancer set regularly serves as the working definition of an “enhancer”for all analyses and individuals.

We evaluated the robustness of this “single definition” approach by performing a comprehensive analysis of similarities in genomic, evolutionary, and functional attributes of enhancers identified by different strategies in two tissues (liver and heart) and two cell lines (K562 and Gm12878). By comparing characteristics of different enhancer sets identified in the same biological context, we were able to assess the stability of conclusions made using only one enhancer identification strategy. While we expected some variation due to differences in the underlying assays, we found significant differences between enhancer sets identified in the same context. These differences between identification strategies are substantial enough to influence biological interpretations and conclusions about enhancer evolution and disease-associated variant function. In general, enhancers defined by cap analysis of gene expression are modestly more enriched for proxies of function than those identified based on evolutionary or functional genomics data; however, they identify a much smaller fraction of active enhancers than other approaches. Additionally, we demonstrate that focusing on enhancers supported by multiple identification methods does not resolve this challenge and ignores many functional enhancers. As a result, we cannot recommend the use of any single current enhancer identification strategy in isolation or simply considering the intersection of many strategies. These results highlight a fundamental challenge to studying gene regulatory mechanisms, and to evaluating the functional relevance of thousands of non-coding variants associated with traits, for instance from GWAS. Incorporating the unique characteristics of different enhancer identification strategies will be essential to ensuring reproducible results, and to furthering our understanding of enhancer biology. As a step toward this goal, we have created creDB, a large, easily queried database of enhancers identified by different methods across common tissues and cellular contexts.

## RESULTS

### A panel of enhancer identification strategies across four biological contexts

To evaluate the variation in enhancer sets annotated by different enhancer identification strategies, we developed a consistent computational pipeline to compare enhancer sets genome-wide. Our approach is based on publicly available data, and we applied it to a representative set of methods in four common cell types and tissues (biological contexts): K562, Gm12878, liver, and heart cells (Figure 1 and Methods). Given the large number of enhancer identification strategies that have been proposed^1,11^, it is not possible to compare them all; so for each context, we considered methods that represent the diversity of experimental and computational approaches in common use.

**Figure 1.**
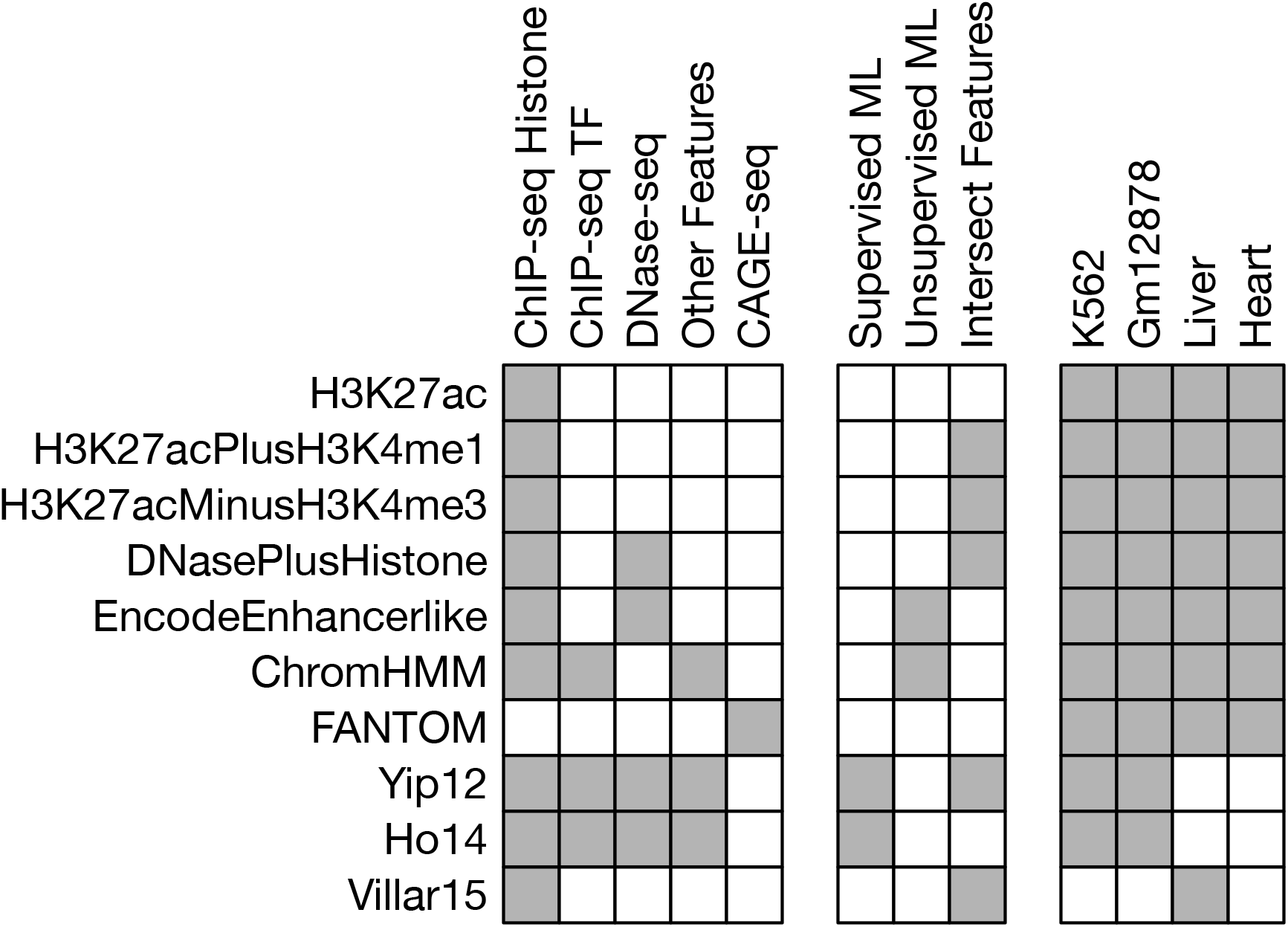
Ten diverse enhancer identification strategies were evaluated across four cellular contexts. Each row summarizes the data sources, analytical approaches, and contexts for the ten enhancer identification strategies we considered. The leftmost columns of the schematic represent the experimental assays and sources of the data used by each identification strategy. The middle columns describe the computational processing (if any) performed on the raw data (ML: machine learning). The rightmost columns give the contexts in which the sets were available. Table 1 gives the number, length, and genomic coverage of each enhancer set.

For all contexts, we considered three enhancer sets derived solely from chromatin immunoprecipitation followed by sequencing (ChIP-seq) for histone modifications informative about enhancer activity from the ENCODE Project^33^. The “H3K27ac” set includes all H3K27ac ChIP-seq peaks without additional refinement^58–62^. The “H3K27acPlusH3K4me1” set includes all H3K27ac ChIP-seq peaks that also overlap an H3K4me1 peak, and the “H3K27acMinusH3K4me3” set contains H3K27ac peaks that do not overlap an H3K4me3 peak^38,39,63^. In liver only, we have an additional set of enhancers identified from H3K27ac ChIP-seq peaks without overlapping H3K4me3 peaks for comparison (“Villar15”)^38^. We also considered a method that incorporates DNase I hypersensitive sites (DHSs) with histone modifications to generate the “DNasePlusHistone” enhancer set, which is composed of DHSs where the ratio of H3K4me1 to H3K4me3 is less than one^40^. Since transcriptional signatures are increasingly used to identify enhancers, we consider “FANTOM” enhancers identified from bidirectionally transcribed eRNA detected via cap analysis of gene expression (CAGE) by the FANTOM5 Project^26,64,65^. We also include several methods that combine machine learning with functional genomics data, such as the ENCODE consortium’s “EncodeEnhancerlike” made by combining DHSs and H3K27ac peaks using an unsupervised ranking method and the “ChromHMM” predictions generated by a hidden Markov model trained on ChIP-seq data from eight histone modifications, CTCF, and RNA Pol II^15,66–68^. For the K562 and Gm12878 cell lines we also include enhancer predictions made by two supervised machine learning methods trained to identify enhancers based on ChIP-seq data in conjunction with other functional genomic features. We will refer to these sets as “Yip12” and “Ho14”^36,37^. An overview of the data and computational approaches used by each method is given in Figure 1 and full details are available in the Methods.

### Enhancers identified by different strategies in the same context differ substantially

#### The genomic coverage of different enhancer sets varies by several orders of magnitude

Enhancer regions identified in the same context by different methods differ drastically in the number of enhancers identified, their genomic locations, their lengths, and their coverage of the genome (Table 1; Figure 2; Figure S1). Different identification methods assay different aspects of enhancer biology (e.g., co-factor binding, histone modification, enhancer RNAs), and therefore we expected to find variation among enhancer sets. Nevertheless, the magnitude of differences we observed is striking. For each attribute we considered, enhancer sets differ by several orders of magnitude (Table 1; Figure 2). For instance, FANTOM identifies 326 kilobases (kb) of sequence with liver enhancer activity, EncodeEnhancerlike identifies 89 megabases (Mb), and H3K27acMinusH3K4me3 identifies almost 138 megabases (Mb). In addition, methods based on similar approaches often differ substantially; e.g., Villar15, which uses the same enhancer definition as H3K27acMinusH3K4me3, only annotates 86.1 Mb with enhancer function in liver. Enhancer sets also vary in their relative distance to other functional genomic features, such as transcription start sites (TSSs). For example, in liver, the average distance to the nearest TSS ranges from 14 kb for EncodeEnhancerlike to 64 kb for DNasePlusHistone (Table S1). Overall, methods based on histone modifications tend to identify larger numbers of longer enhancers compared with CAGE data, and machine learning methods are variable. We highlight these trends in liver, but they are similar in other contexts (Table 1; Figure 2; Figure S1).

**Figure 2.**
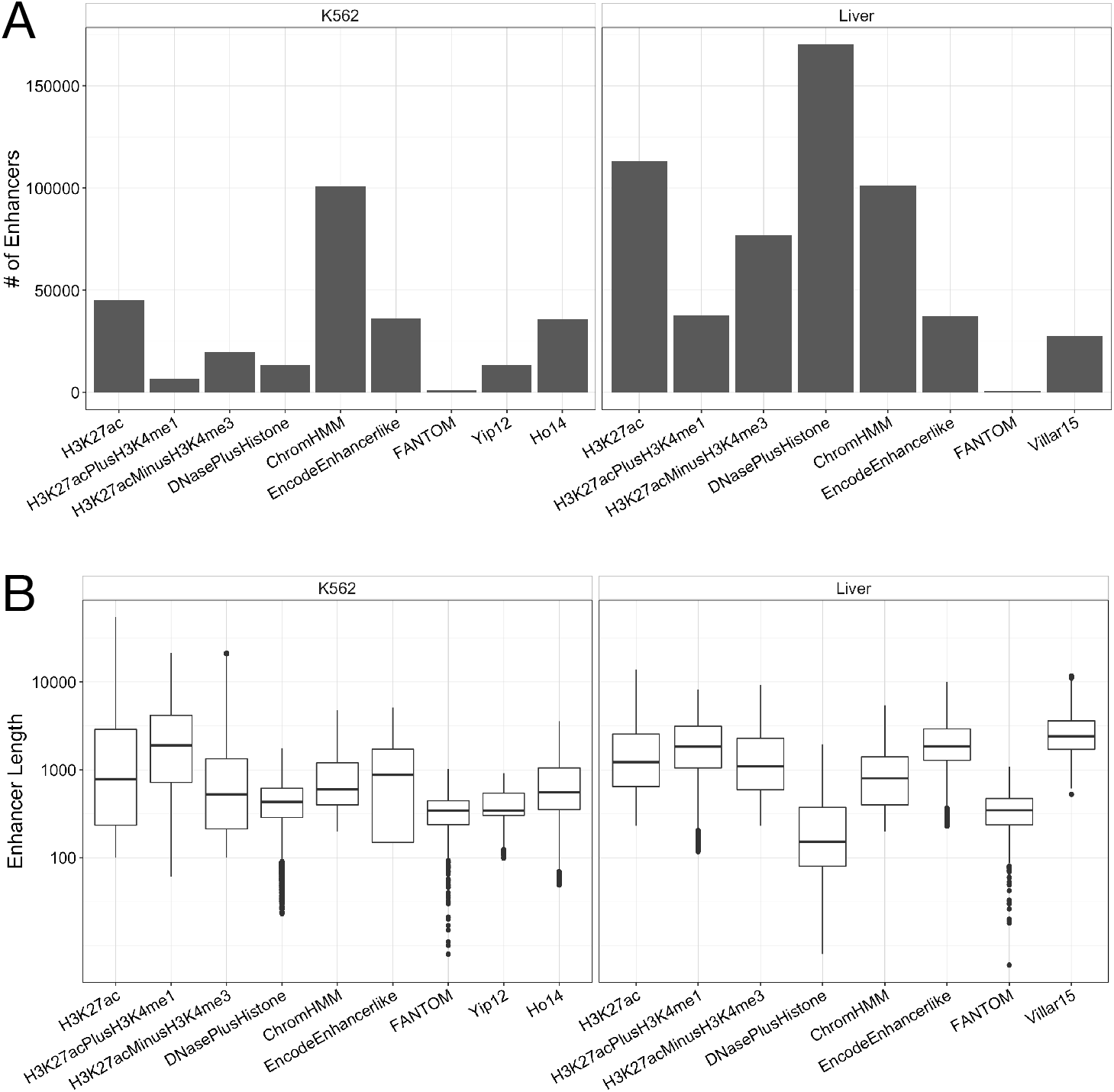
Enhancer identification methods vary in the number and length of predicted enhancers. (A) The number of K562 and liver enhancers identified by each method varies over two orders of magnitude. There is considerable variation even among methods defined based on similar input data, e.g., histone modifications. (B) The length of K562 and liver enhancers identified by different methods shows similar variation. Enhancer lengths are plotted on a log10 scale on the y-axis. Data for other contexts are available in Table 1 and Figure S1.

**Table 1.**
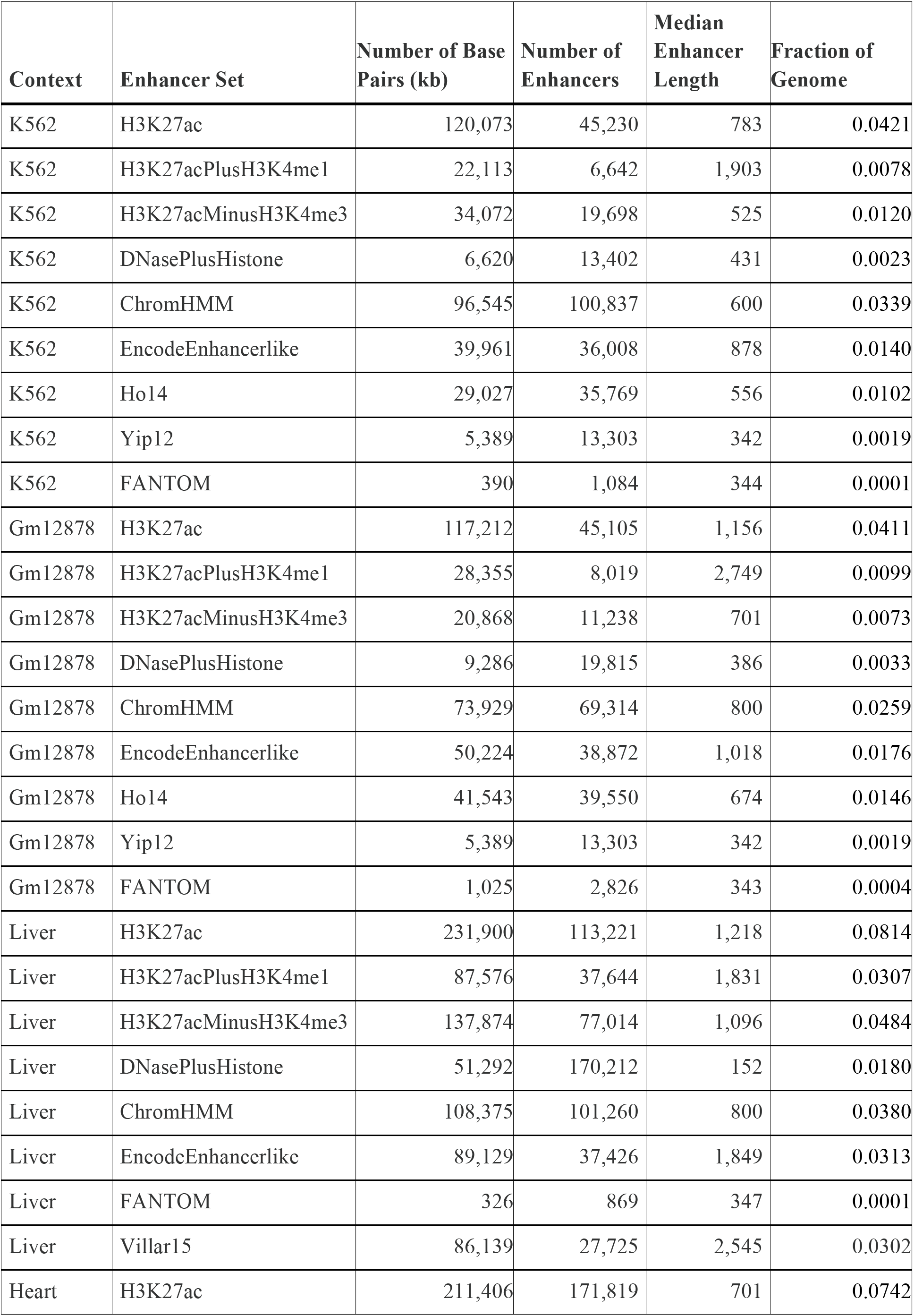
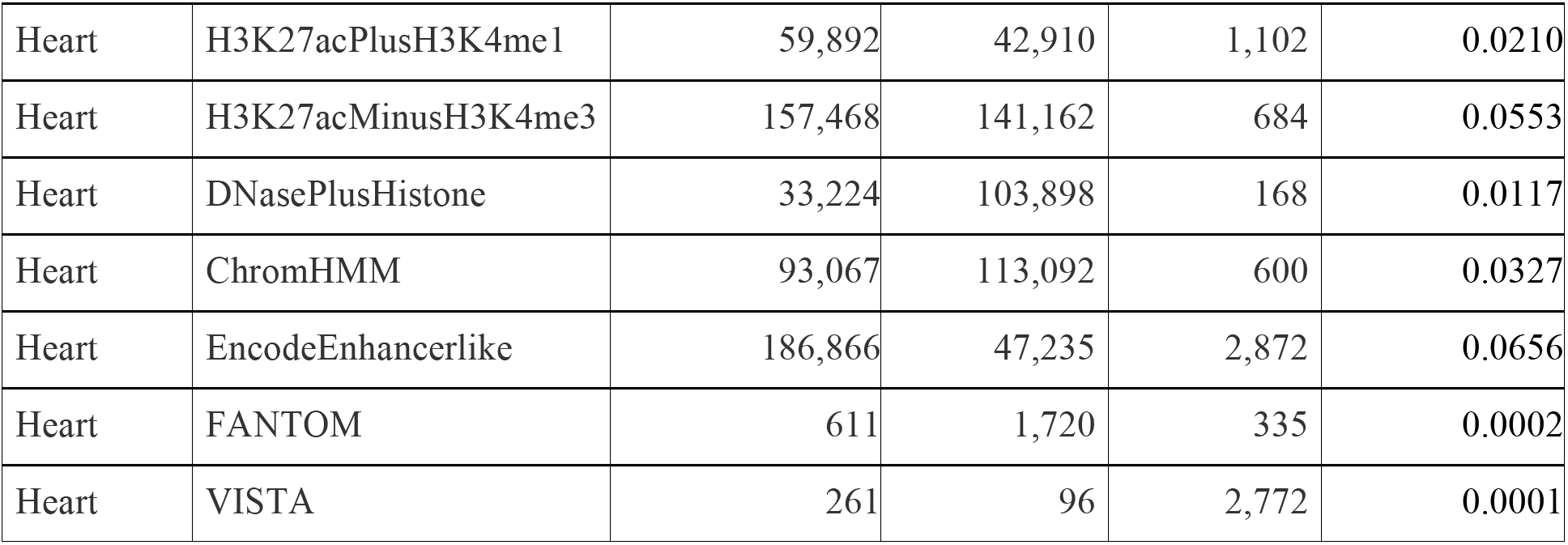
Summary of all enhancer sets analyzed in this study.

#### Enhancer sets overlap more than expected by chance, but have low genomic similarity

Given the diversity of the enhancer sets identified by different methods, we evaluated the extent of bp overlap between them. All pairs of enhancer sets overlap more than one could expect if they were randomly distributed across the genome (Figure 3A,B, p < 0.001 for all pairs). As expected due to the greater cellular heterogeneity and genetic variation of tissue samples vs. cell lines, enhancer sets identified by different methods in the same cell line have more significant overlap than enhancer sets identified in tissues (Figure 3B). However, the magnitude of overlap between enhancer sets is typically low: less than 50% for nearly all pairs of methods (median 20% bp overlap for K562 and 31% for liver; Figure 3C,D, Figure S2A,B; Table S2). Indeed, most (54% across contexts) annotated regions are “singletons” that are annotated by only a single enhancer identification strategy. Furthermore, the largest overlaps are in comparisons including one enhancer set with high genome coverage, or in comparisons of sets that were identified based on similar data. These patterns were similar when evaluating overlap on an element-wise basis (Figures S3-S4, Table S2), as opposed to single bp resolution. To further quantify overlap, we calculated the Jaccard similarity index—the number of shared bp between two enhancer sets divided by the number of bp in their union—for each pair of methods. Overall, the Jaccard similarities are also extremely low for all contexts, with an average of 0.12 for K562 and 0.17 for liver and the majority of pairwise comparisons below 0.30 (Figure 3E,F, Figure S2C,D, upper triangle). Since the Jaccard similarity is sensitive to differences in set size, we also computed a “relative” Jaccard similarity by dividing the observed value by the maximum value possible given the set sizes. The relative similarities were also consistently low (Figure 3E,F, Figure S2C,D, lower triangle).

**Figure 3.**
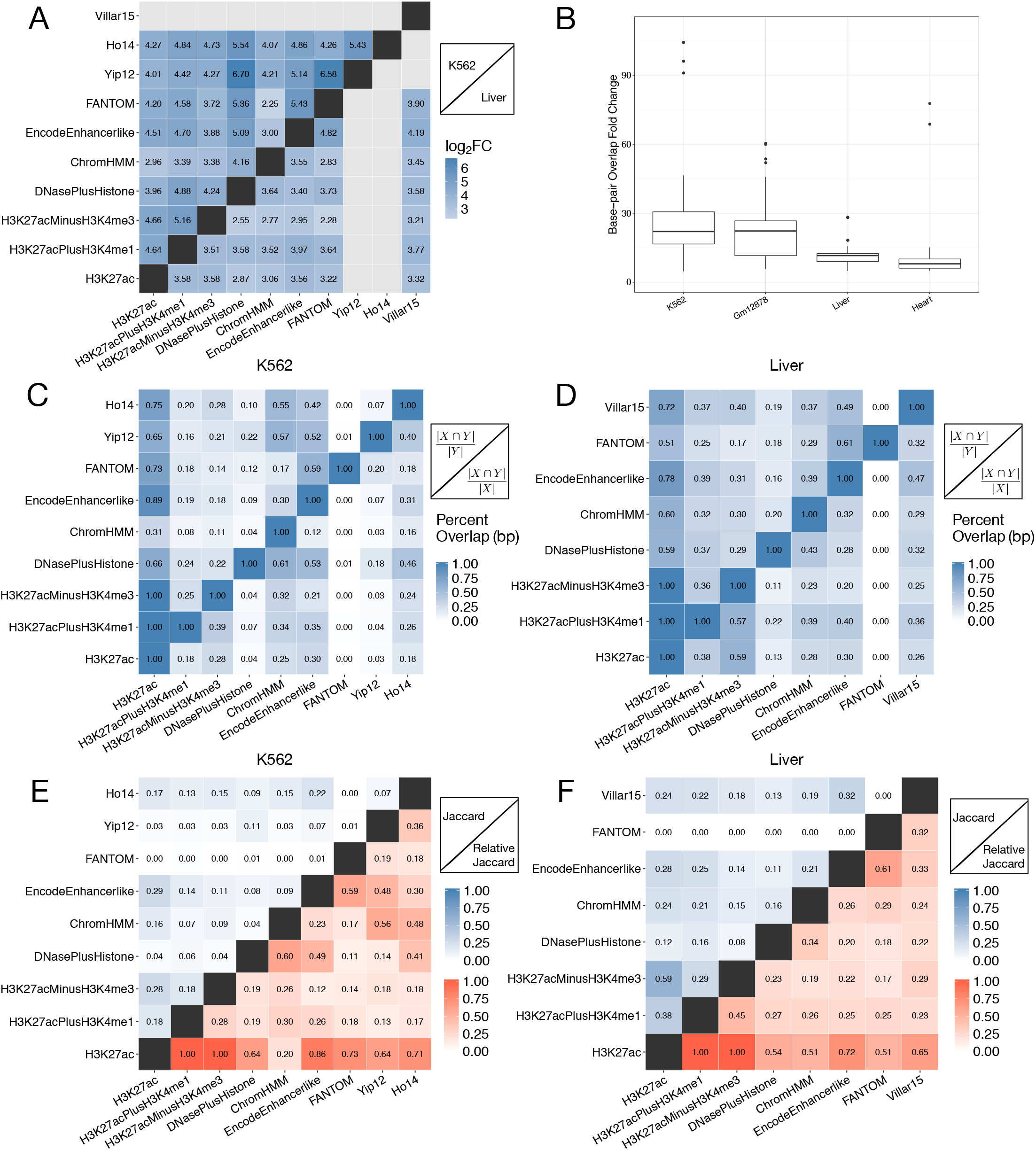
Enhancer sets have low genomic overlap. (A) Pairwise bp enrichment values (log_2_ fold change) for overlap between each K562 (upper triangle) or liver (lower triangle) enhancer set, compared to the expected overlap between randomly distributed, length-matched regions. (B) The enrichment for bp overlap compared to a random genomic distribution for each pair of enhancer sets within each context. The fold changes for the primary tissues—liver and heart—are significantly lower than the cell lines—K562 and Gm12878 (p = 4.11E-21 Kruskal-Wallis test, followed by Dunn’s test with Bonferroni correction for pairwise comparisons between contexts). The patterns are similar for element-wise comparisons (Figures S3 and S4). (C, D) The percent base pair (bp) overlap between all pairs of (C) K562 enhancer sets and (D) liver enhancer sets. Percent overlap for each pair was calculated by dividing the number of shared bp between the two sets by the total number of base pairs of the set on the y-axis. High overlap is observed only for pairs based on very similar data, e.g., H3K27ac, H3K27acPlusH3K4me1, and H3K27acMinusH3K4me3, or comparisons with the largest set, e.g. H3K27ac. (E, F) The Jaccard similarity between all pairs of (E) K562 or (F) liver enhancer sets. The upper triangle gives the Jaccard similarity, and the lower triangle gives the relative Jaccard similarity in which the observed similarity is divided by the maximum possible similarity for the pair of sets.

To assess the influence of technical variation on the observed overlaps, we compared the overlap of replicates from H3K27ac ChIP-seq data in the K562 and Gm12878 cell lines generated by the same laboratory. We expected the replicates to have high overlap and serve as an “upper bound” on similarity in practical applications. The replicates overlap 65–80% at the bp level. Thus, while there is variation, the amount of overlap observed between enhancers identified by different methods almost always falls far below the variation between ChIP-seq replicates.

#### Enhancer sets have different levels of evolutionary conservation

Enhancers identified by different methods also differ in their levels of evolutionary constraint. Using primate and vertebrate evolutionarily conserved elements defined by PhastCons^42^, we calculated the enrichment for overlap with conserved elements for each enhancer set. All enhancer sets have more regions that overlap with conserved elements than expected from length-matched regions drawn at random from the genome (Methods). However, enhancers identified by some methods are more likely to be conserved than others (Figure 4A). Across each context, the histone-based, ChromHMM, Villar15, and Ho14 enhancer sets are approximately 1.3x to 1.8x enriched for overlap with conserved elements. Adding DNaseI hypersensitivity data, as in the DNasePlusHistone and EncodeEnhancerlike sets, increases the level of enrichment slightly compared to solely histone-derived enhancers (1.9x–2.3x). In contrast, the FANTOM and Yip12 enhancers are nearly twice as enriched for conserved regions as the histone-based sets (2.7x and 3.3x, respectively). Evolutionary conservation was not directly considered in the definition of the FANTOM and Yip12 sets. Here we considered enhancer elements overlapped by conserved elements; enrichment trends are similar when we consider the number of conserved base pairs overlapped by each enhancer set (Figure S5A).

**Figure 4.**
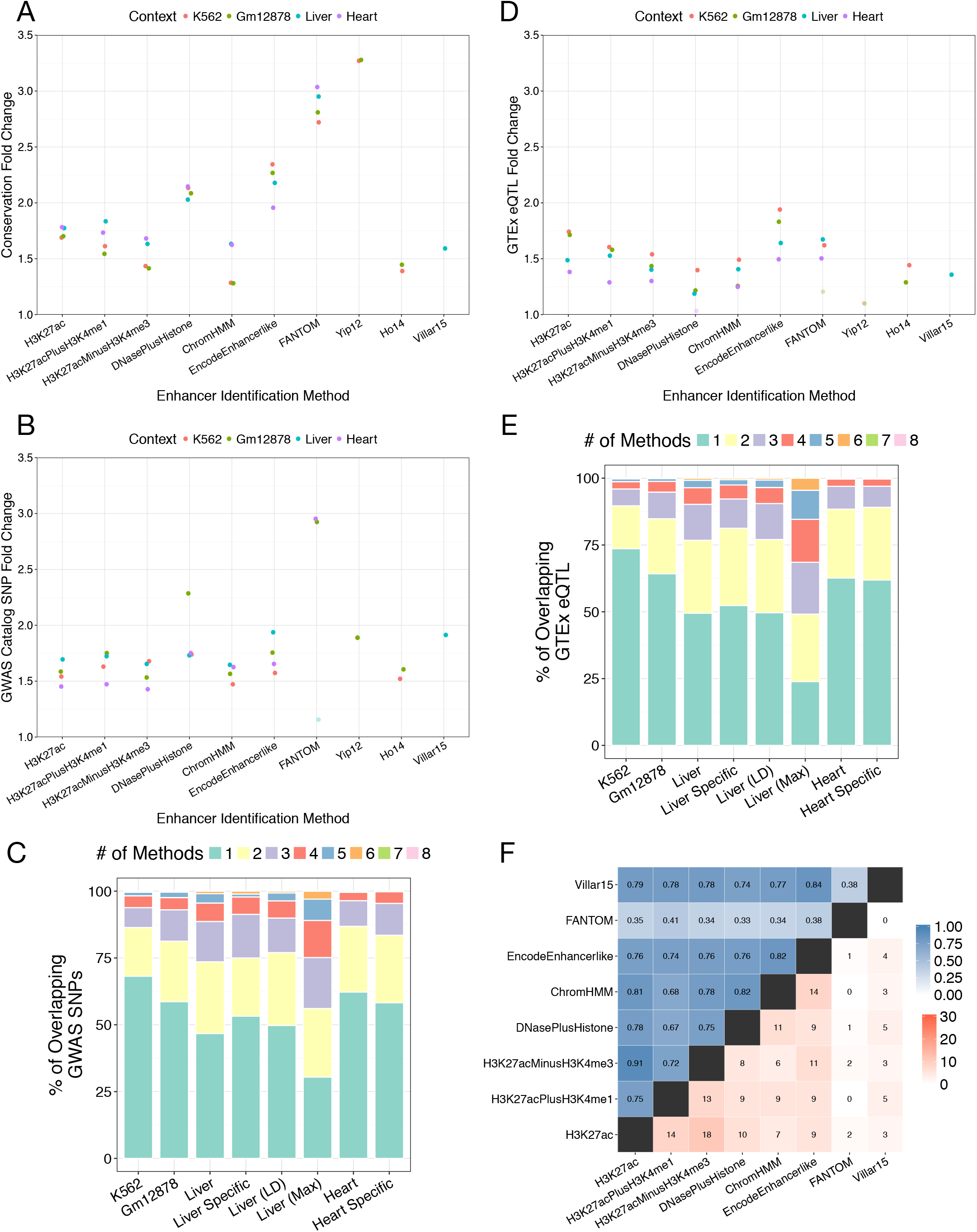
Enhancers identified by different methods differ in functional attributes. (A) Enhancer sets vary in their degree of evolutionary conservation. Each point represents the enrichment (fold change compared to randomly shuffled regions) for overlap between a conserved element (combined primate and vertebrate PhastCons) and each enhancer set. FANTOM and Yip12 are the most enriched for conserved elements, while sets based on histone modification data alone are among the least enriched (B) GWAS SNP enrichment among all enhancer sets for each biological context. All sets are significantly enriched, except FANTOM in K562 and liver contexts due to small sample size. (C) Among all GWAS SNPs that overlap at least one enhancer in a context, the colored bars represent the number of methods that identified the region as an enhancer. The majority of these variants are supported by a single method; very few GWAS variants are shared among all methods. The conclusions are similar when considering variants in high LD (r^2^ > 0.9) with the GWAS tag SNPs in liver (Liver LD; Figure S8). The pattern is also similar when limiting to SNPs associated with liver or heart related phenotypes (Liver Specific, Heart Specific). When considering the SNP in each LD block with the maximum number of enhancer overlaps there is still a large percentage of SNPs supported by only one method (Liver Max). (D) GTEx eQTL enrichment among all enhancer sets for each biological context. All sets were enriched except for DNasePlusHistone in heart. (E) Among all eQTL that overlap at least one enhancer, the majority is supported by only a single method. This holds for LD-expanded and context-specific sets (Liver LD, Liver Specific, Heart Specific; Figure S8). Many variants remain unique to a single method, even when limiting to the variant in each LD block overlapping the maximum of enhancer sets (Liver Max). These trends are similar to what is seen for GWAS SNPs in (C). (F) Enhancer sets from the same biological context have different functional associations. We identified Gene Ontology (GO) functional annotations enriched among genes likely to be regulated by each enhancer set using GREAT. The upper triangle represents the pairwise semantic similarity for significant molecular function (MF) GO terms associated with predicted liver enhancers. The lower triangle shows the number of shared MF GO terms in the top 30 significant hits for liver enhancer sets. Results were similar when using enhancer-gene target predictions from JEME (Figures S9 and S10).

#### Identification strategies highlight different subsets of experimentally validated enhancers

Though we lack unbiased genome-wide gold-standard sets of enhancers, nearly two thousand human sequences have been tested for enhancer activity *in vivo* in transgenic mice at E11.5 by VISTA^46^ and thousands more have been tested in cell lines via massively parallel reporter assays (MPRAs). Strong ascertainment biases in how regions were selected for testing in these assays prevent their use as a gold standard, but they do provide an opportunity to examine overlap between validated and predicted enhancers. We evaluated the overlap and enrichment of each heart enhancer set with 1,837 regions tested for enhancer activity in the developing heart by VISTA and for each annotated K562 enhancer with 15,720 regions tested in K562 cells by Sharpr-MPRA^47^. All heart enhancer sets are significantly enriched for overlap with the 126 VISTA heart positives (Figure S6A; p < 0.001 for all), and each set is at least ~3x more likely to overlap validated enhancers than expected if it was randomly distributed across the genome. However, the heart enhancer sets are also significantly enriched for overlap with VISTA negatives (p <= 0.004). This is not surprising as the regions tested by VISTA were largely selected based on having evidence of enhancer activity, and they may have enhancer activity in other contexts not tested by VISTA, including adult heart. Furthermore, there is substantial disagreement among the enhancer sets about the status of the VISTA heart enhancers; 16% (n = 20) of validated heart enhancers are not predicted to have enhancer activity by any method, and 17% (22) are only predicted by one method (Figure S6B,C). Similarly, all of the enhancer sets in K562 are significantly enriched for overlap with both activating and repressive regions characterized by Sharpr-MPRA (Figure S7A; p < 0.001 for all). There is little variation between the methods in terms of overall enrichment, with most being ~2x enriched for activating regions. Nearly half of the activating regions in the MPRA (49%; 2,615 / 5,373) were not identified by any of the enhancer sets, and 40% of activating regions overlapping a predicted enhancer are unique to a single set (Figure S7B,C; 1,098 / 2,747). Thus, comparison with validated enhancers from both VISTA and MPRAs suggest that different strategies identify different subsets of active regulatory regions in the same context, and that all strategies miss a sizable portion of functional enhancer sequences. However, we again caution against interpreting the relative performance of different enhancer identification strategies on these data, since there are strong ascertainment biases in how regions were selected for testing. For example, ChromHMM enhancer predictions and DNase I hypersensitivity data were used to select the regions tested by Sharpr-MPRA.

### Interpretation of GWAS hits and eQTL is contingent on the enhancer identification strategy used

Genome-wide enhancer sets are commonly used to interpret the potential function of genetic variants observed in GWAS and sequencing studies^62,63,65,68–74^. Functional genetic variants—in particular mutations associated with complex disease— are enriched in gene regulatory regions^5,6^. We evaluated the sensitivity of this pattern to the enhancer identification strategy used by intersecting each of the enhancer sets with 20,458 unique tag SNPs significantly associated with traits from the NHGRI-EBI GWAS Catalog. Overall, 34.9% (7,133 / 20,458) of GWAS SNPs overlap an enhancer identified by at least one of the strategies in one of the contexts we considered. However, there is wide variation in the number of overlapping GWAS Catalog SNPs between enhancer sets, as is expected given the large variation in the number and genomic distribution of enhancers predicted by different methods (Table S4). Nonetheless, GWAS tag SNPs are significantly enriched at similar levels in most enhancer sets and contexts, with the exception of FANTOM, which has roughly twice the enrichment of other methods in Gm12878 and heart (Figure 4B). Since the tag SNPs are often not the functional variants, we also considered SNPs in high linkage disequilibrium (LD) with the GWAS SNPs (r^2^ > 0.9). While the overall enrichments were lower, the variability between enhancer sets remained small (Figure S8A). We also identified the variant in each LD block with the maximum number of overlaps with distinct enhancer sets. Even when limiting our analysis to this biased set, 58% of GWAS SNPs (11,854 / 20,458) have no enhancer overlap (Figure S8B).

Furthermore, GWAS SNPs with enhancer overlap are commonly predicted to overlap an enhancer by only a single identification strategy (Figure 4C). For example, in liver, 47% (1710 / 3660) of the GWAS SNPs that overlapped an enhancer are unique to a single set, and only 26% (968 / 3660) overlap enhancers from more than two sets. The distribution of enhancer overlaps was similar when considering all candidate variants in LD (Figure 4C). Even after limiting to GWAS LD blocks with enhancer overlap and selecting the variant with maximum overlap, 30% (2620 / 8604) still are only predicted by one enhancer identification method (Figure 4C). This illustrates that the annotation of variants in regions highlighted by GWAS varies greatly depending on the enhancer identification strategies used. Since the GWAS catalog contains regions associated with diverse traits, we manually curated the set of GWAS SNPs into subsets associated with phenotypes relevant to liver or heart (Table S5). As in the full GWAS set, the majority of curated GWAS liver SNPs with any enhancer overlap are overlapped by a single method (53%) and none are shared by all methods (Figure 4C). The heart and liver enhancer sets are almost universally more enriched for overlap with GWAS SNPs that influence relevant phenotypes compared to GWAS SNPs overall (Table S6; 1.74x–2.68x). FANTOM enhancers are the exception to this trend due to the small number of overlapping context-specific SNPs (Table S4). This suggests that the different methods, in spite of their lack of agreement, all identify regulatory regions relevant to the target context.

To test if these patterns hold for genetic variants in other functional regions, we analyzed the overlap of enhancer sets with expression quantitative trait loci (eQTL) identified by the GTEx Consortium. These analyses yielded similar results as for the GWAS Catalog variants (Figure 4D). Within a context, most eQTL are identified as enhancers by a single enhancer prediction method only, and there is wide variation in the number and enrichment of eQTL overlapped by different enhancer sets (Figure 4D; Table S7). Across liver enhancer sets, 50% (33,941 / 68,563) of all overlapped eQTL are called an enhancer by only a single method (Figure 4E). Considering variants in high LD (r^2^ > 0.9) does not affect this trend (Figure 4E). Similarly, after limiting the analysis to the variants with the maximum number of overlaps in each LD block, 24% (64871 / 271732) of the eQTL with enhancer overlap are identified by only one enhancer set (Figure 4E). Furthermore, restriction to context-specific eQTL in liver or heart does increase the enrichment for eQTL across most methods, but the distribution of shared eQTL remains similar (Figure 4E; Table S8). Thus, the interpretation of variants in regions known to influence gene regulation varies substantially depending on the enhancer identification strategy used.

### Enhancers identified by different strategies have different functional contexts

Given the genomic dissimilarities between enhancer sets, we hypothesized that different enhancer sets from the same context would also vary in the functions of the genes they regulate. To test this hypothesis, we identified Gene Ontology (GO) functional annotation terms that are significantly enriched among genes likely targeted by enhancers in each set. We used two different approaches to discover genes and associated GO terms: (*i*) using the joint effect of multiple enhancers (JEME) method for mapping enhancers to putative target genes and then performing enrichment analyses, and (*ii*) applying the Genomic Regions Enrichment of Annotations Tool (GREAT) (Methods)^51,54^. Many of the GO terms identified by both methods for the enhancer sets are relevant to the associated context (Table 2). However, most of the associated terms for the target-mapping approach were near the root of the ontologies and thus lacking in functional specificity (Table 2), likely due to the large gene target lists for most enhancer sets (Table S9). As a result, we focus on the results from GREAT here, and report the results based on JEME target mapping in Figures S9 and S10.

**Table 2.** Top 5 Gene Ontology (Molecular Function) terms for liver enhancer sets from GREAT and JEME target-mapped WebGestalt enrichments.

| Enhancer Set | GO MF Terms (GREAT) | GO MF Terms (JEME+WebGestalt) |
| --- | --- | --- |
| H3K27ac | cytoskeletal adaptor activity | nucleotide binding |
|  | 14-3-3 protein binding | nucleoside binding |
|  | nucleobase-containing compound transmembrane transporter activity | purine nucleoside binding |
|  | high-density lipoprotein particle binding | DNA binding |
|  | JUN kinase binding | RNA binding |
| H3K27acPlusH3K4me1 | cytoskeletal adaptor activity | small molecule binding |
|  | 14-3-3 protein binding | anion binding |
|  | leukotriene-C4 synthase activity | nucleoside phosphate binding |
|  | nucleobase-containing compound transmembrane transporter activity | nucleotide binding |
|  | FAD binding | transferase activity |
| H3K27acMinusH3K4me3 | 14-3-3 protein binding | oxidoreductase activity |
|  | cytoskeletal adaptor activity | anion binding |
|  | thyroid hormone receptor binding | small molecule binding |
|  | ARF guanyl-nucleotide exchange factor activity | nucleoside phosphate binding |
|  | high-density lipoprotein particle binding | nucleotide binding |
| DNasePlusHistone | cytoskeletal adaptor activity | small molecule binding |
|  | glucocorticoid receptor binding | anion binding |
|  | nucleobase-containing compound transmembrane transporter activity | transferase activity |
|  | high-density lipoprotein particle binding | nucleotide binding |
|  | 14-3-3- protein binding | nucleoside phosphate binding |
| ChromHMM | high-density lipoprotein particle binding | nucleotide binding |
|  | nucleobase-containing compound transmembrane transporter activity | nucleoside binding |
|  | cytoskeletal adaptor activity | purine nucleoside binding |
|  | 14-3-3 protein binding | DNA binding |
|  | retinoid X receptor binding | RNA binding |
| EncodeEnhancerlike | cytoskeletal adaptor activity | nucleotide binding |
|  | 14-3-3 protein binding | transferase activity |
|  | nucleobase-containing compound transmembrane transporter activity | small molecule binding |
|  | apolipoprotein A-I binding | anion binding |
|  | high-density lipoprotein particle binding | carbohydrate derivative binding |
| FANTOM | glucocorticoid receptor binding | structural constituent of ribosome |
|  | protein kinase binding | receptor binding |
|  | kinase binding | cell adhesion molecule binding |
|  | methylglutaconyl-CoA hydratase activity | molecular function regulator |
|  | vitamin D response element binding | transcription regulatory region DNA binding |
| Villar15 | protease binding | anion binding |
|  | phosphatidylinositol 3-kinase binding | small molecule binding |
|  | 14-3-3 protein binding | oxidoreductase activity |
|  | cytoskeletal adaptor activity | cofactor binding |
|  | glucocorticoid receptor binding | oxidoreductase activity, acting on CH-OH group of donors |

The majority of the top 30 significant annotations from GREAT for each enhancer set are not enriched in any other set in the same context, and no terms are shared by all of the methods in a given context (Figure 4F, lower triangle). In most of these pairwise comparisons, fewer than half of the GO terms are shared between a pair of enhancer sets. Furthermore, many of the terms shared by multiple enhancer sets are near the root of the ontology and thus are less functionally specific. These results provide evidence that the different enhancer sets influence different functions relevant to the target biological context. These trends hold for both the Biological Process (BP) and Molecular Function (MF) ontologies and considering the top 10 and 50 annotations for each set (Figures S11-S13).

To further compare the enriched GO MF and BP annotations of each enhancer set in a way that accounts for the distance between GO terms in the ontology hierarchy and their specificity, we computed a semantic similarity measure developed for GO annotations^52,53^. The histone-derived H3K27ac and H3K27acMinusH3K4me3 enhancer sets are among the most functionally similar, with similarity scores of at least 0.80 in most contexts (Figures 4F, upper triangle; Figure S12). This is not surprising given that their underlying assays overlap. The functional similarity scores are lower for comparisons of the other histone modification sets, around 0.50–0.80. In all comparisons, the FANTOM enhancers have the lowest functional similarity with other enhancer sets—below 0.40 in the vast majority of comparisons in K562, liver, and heart (Figure 4F; Figure S12). FANTOM is more similar to other methods in Gm12878, with an average score of 0.59 (Figure S12). As a benchmark, biological replicates for the Gm12878 H3K27ac enhancer set received a similarity of 0.93. This suggests different functional influences for enhancer sets from the same context identified by different methods, with FANTOM as a particular outlier. We note that enhancer target gene identification remains a challenging problem, and both strategies for mapping enhancers to potential target genes considered here (GREAT and JEME) likely include false positives. However, insofar as they reflect the regulatory context of the different enhancer sets, they reveal significant functional differences between enhancer identification methods.

### Genomic and functional clustering of enhancer sets

Our analyses of enhancer sets within the same biological context reveal widespread dissimilarity in both genomic and functional features. To summarize and compare the overall genomic and functional similarity of the enhancer sets across contexts, we clustered them using hierarchical clustering and multidimensional scaling (MDS) based on their Jaccard similarity in genomic space and the GO term functional similarity of predicted target genes (Methods).

Several trends emerged from analyzing the genomic and functional distribution within and between biological contexts. First, the FANTOM eRNA enhancers are consistently distinct from all other enhancer sets in both their genomic distribution and functional associations (Figure 5). Differences between eRNA and non-eRNA enhancer sets appear to dominate any other variation introduced by biological, technical, or methodological differences. Second, similarity in genomic distribution of enhancer sets does not necessarily translate to similarity in functional space, and vice versa. For example, although DNasePlusHistone enhancers are close to the other histone-derived enhancer sets and the machine learning models in the functional comparisons (Figure 5B, D), they are located far from those sets in the genomic-location-based projection and hierarchical clustering (Figure 5A, C). Histone-derived enhancer sets are also farther from the machine-learning-based sets in genomic comparisons than in functional analyses. Finally, comparing enhancer sets by performing hierarchical clustering within and between biological contexts reveals that genomic distributions are generally more similar within biological contexts, compared to other sets defined by the same method in a different context (Figure 5E). For example, the ChromHMM set from heart is more similar to other heart enhancer sets than to ChromHMM sets from other contexts. In contrast, the enhancer set similarities in functional space are less conserved by biological context (Figure 5F). Here, the heart ChromHMM set is functionally more similar to the H3K27acMinusH3K4me3 set from liver cells than other heart enhancer sets. In general, cell line enhancer sets (red and green) show more functional continuity than heart and liver sets (blue and purple). However, the FANTOM enhancers are exceptions to these trends; FANTOM enhancers from each context form their own cluster based on their genomic distribution, underscoring their uniqueness.

**Figure 5.**
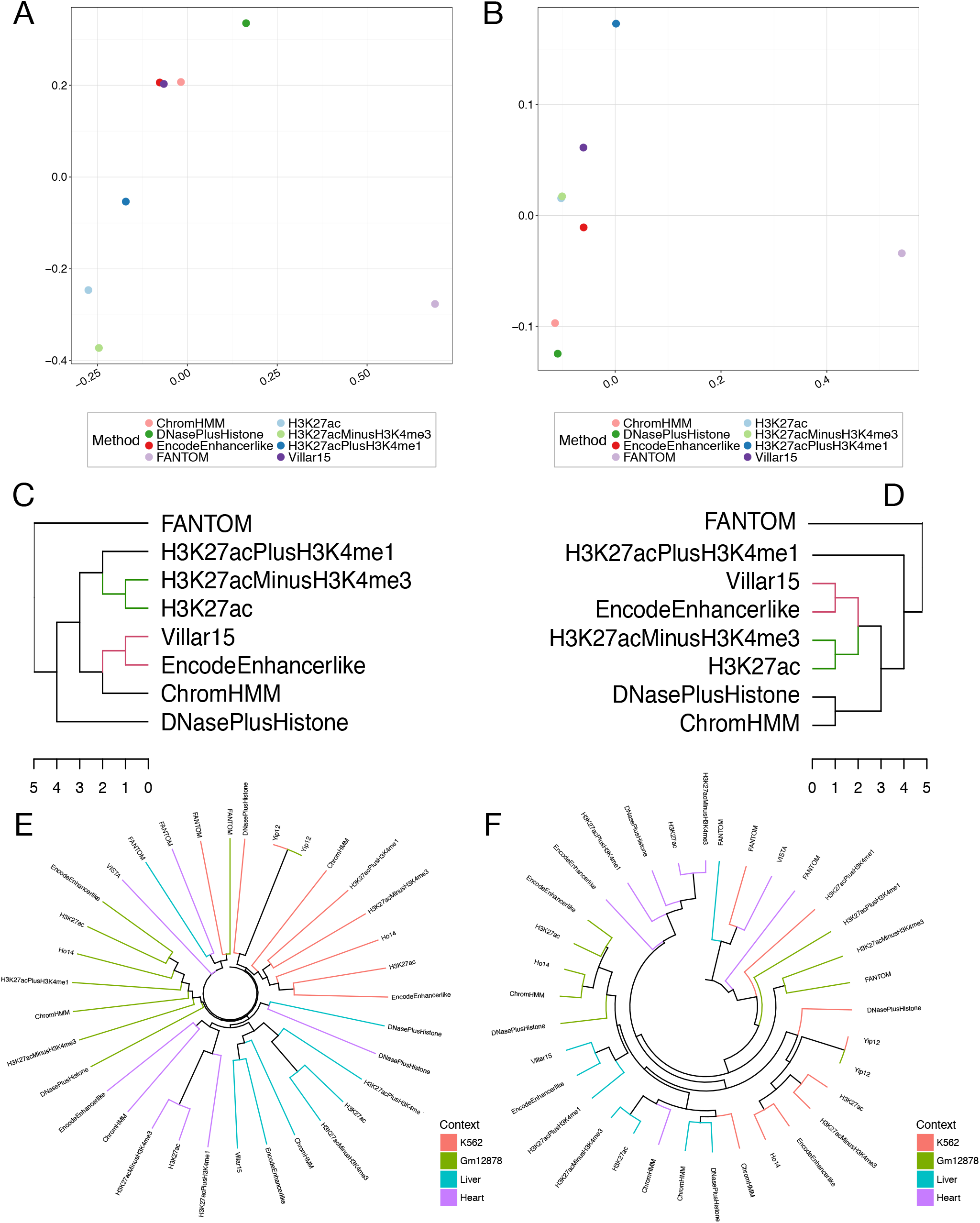
The genomic and functional similarities between enhancer sets are not consistent. (A) Multidimensional scaling (MDS) plot of liver enhancer sets based on the Jaccard similarity of the genomic distributions (Figure 3B). (B) MDS plot for liver enhancers based on distances calculated from molecular function (MF) Gene Ontology (GO) term semantic similarity values (Figure 4F). (C, D) Hierarchical clustering based on the Jaccard similarities of the genomic distributions (C) of all liver enhancer sets compared to clustering based on GO semantic similarity (D). FANTOM enhancers are the most distant from all other enhancer sets in both genomic and functional similarity, but the relationships between other sets are not conserved. Colored branches denote identical subtrees within the hierarchy. (E) Hierarchical clustering based on genomic Jaccard distances for all methods and all contexts. (F) Hierarchical clustering of all available enhancer sets based on GO term distances. Terminal branches in E and F are colored by biological context. With the exception of FANTOM enhancers, the enhancer sets’ genomic distributions are more similar within than between biological contexts. Functional similarity does not always correlate with genomic similarity, and the clustering by biological context is weaker in functional space.

### Combining enhancer sets does not strongly increase evidence for regulatory function

Although there are large discrepancies in genomic and functional attributes between enhancer sets identified by different methods in the same context, we hypothesized that the subset of regions shared by two or more sets would have stronger enrichment for markers of gene regulatory function. To test this, we analyzed whether regions identified by multiple methods have increased “functional support” compared to regions identified by fewer methods. We evaluated three signals of functional importance: *(i)* enrichment for overlap with evolutionarily conserved elements, *(ii)* enrichment for overlap with GWAS SNPs, and *(iii)* enrichment for overlap with GTEx eQTL. For each, there are only small changes as the number of methods identifying a region increases (Figure 6A–C). Regions identified as enhancers by more than one method are slightly more enriched for conserved elements compared to the genomic background, but there is little difference among regions identified by 2–5 methods (Figure 6A). Regions predicted by 6 or more methods are significantly more enriched for conserved elements than those with less support, but effect size is modest (1.36x for 1 vs. 1.62x for 6). There is a modest increase in the enrichment for overlap with GWAS SNPs among enhancers identified by more identification methods; however, given the relatively small number of GWAS SNP overlaps, none of these differences were statistically significant (Figure 6B). We observed no increase in the enrichment for overlap with eQTL as the support for enhancer activity increased (Figure 6C). Thus, we do not find strong evidence of increased functional importance in enhancers identified by multiple methods compared to enhancers identified by a single method. Importantly, this implies that intersecting enhancer identification strategies will focus on a smaller set of enhancers with little evidence for increased functional relevance.

**Figure 6.**
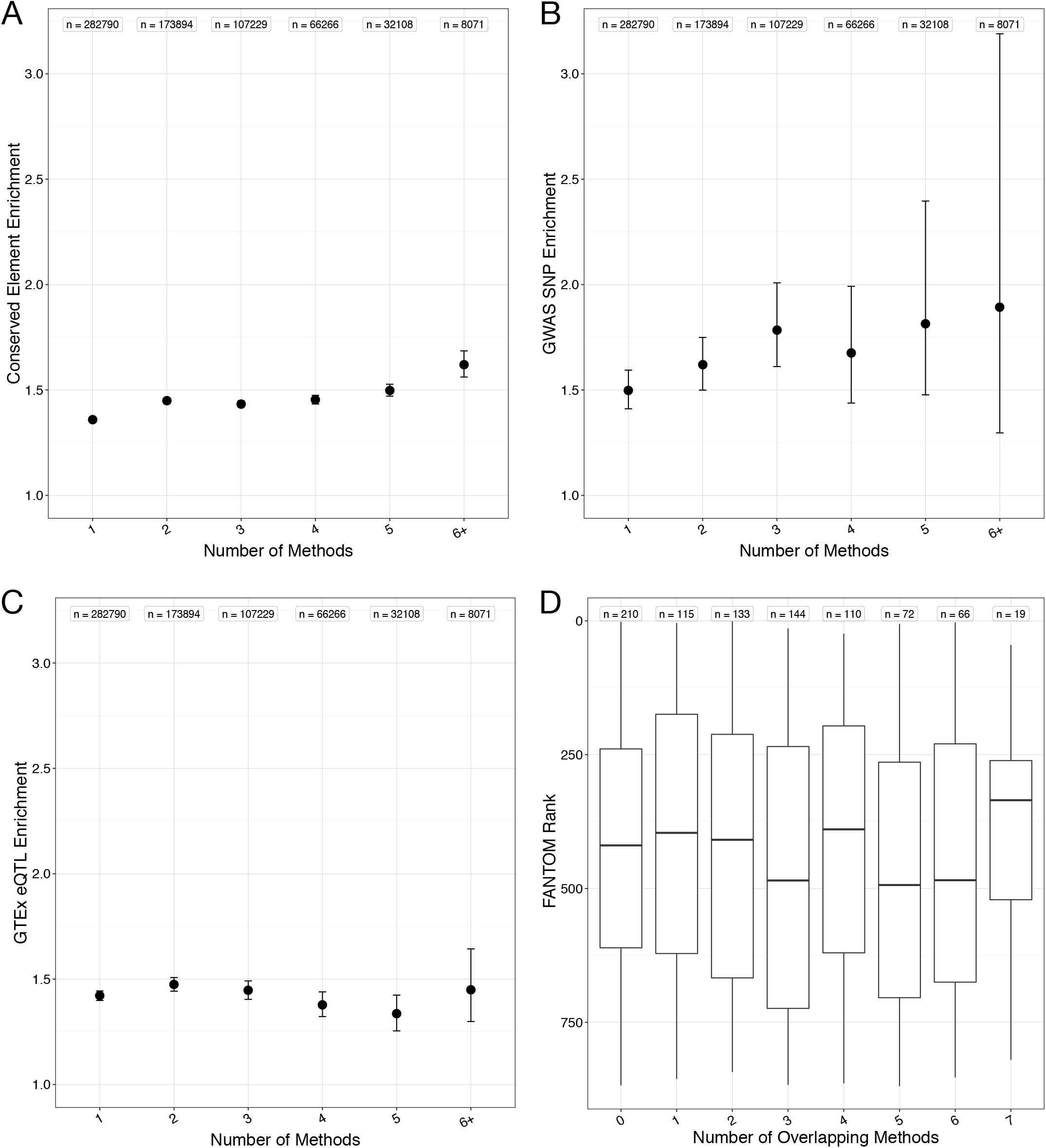
Enhancers identified by multiple methods have little additional evidence of function. (A) Enrichment for overlap between conserved elements (n = 3,930,677) and liver enhancers stratified by the number of identification methods that predicted each enhancer. (B) Enrichment for overlap between GWAS SNPs (n =20,458) and liver enhancers stratified by the number of identification methods that predicted each enhancer. (C) Enrichment for overlap between GTEx eQTL (n = 429,964) and liver enhancers stratified by the number of identification methods that predicted each enhancer. In (A–C), the average enrichment compared to 1,000 random sets is plotted as a circle; error bars represent 95% confidence intervals; and n gives the number of enhancers in each bin. The only significant differences are found in the enrichment for evolutionary conservation (A), but the difference is modest in magnitude (1.36x for 1 vs. 1.62x for 6+). (D) Boxplots showing the distribution of confidence score ranks for FANTOM enhancers in liver partitioned into bins based on the number of other methods that also identify the region as an enhancer. Lower rank indicates higher confidence; note that the y-axis is flipped to the high confidence (low rank) regions are at the top. The lack of increase in enhancer score with the number of methods supporting it held across all methods tested (Figure S14).

Several enhancer identification methods provide confidence scores that reflect the strength of evidence for each enhancer. We hypothesized that high confidence enhancers from one method would be more likely to overlap enhancers identified by other methods. To test this, we ranked each enhancer based on its confidence or signal, with a rank of 1 representing the highest confidence in the set. There was no clear trend between the confidence score of an enhancer from one method and the number of methods that identified the region as an enhancer (Figure 6D, Figure S14-17). Overall, enhancers identified by multiple methods show a similar confidence distribution when compared to regions identified by a single method. Indeed, for some enhancer sets the median score decreases as the regions become more highly shared (Figure S14A–C, Figure S17A–C). This provides further evidence that building enhancer sets by simple combinations of existing methods is unlikely to lead to a higher confidence subset, and that filtering based on simple agreement between methods may not improve the specificity of enhancer predictions.

### A tool for evaluating of the robustness of conclusions across enhancer identification strategies

Our findings imply that results of studies of enhancers are sensitive to the enhancer identification strategy used. To facilitate evaluation of the sensitivity and robustness of results with respect to this choice, we provide an integrated database (creDB) of annotated enhancer sets. It contains all enhancer sets used in this study, and enhancer sets from multiple primary and meta-analyses annotating enhancers across cellular contexts. In contrast to other efforts providing large compilations of enhancer sets^71,72^, creDB is not a web-server or “enhancer browser”. Rather, it is an SQL database with a convenient interface to the R programming language and distributed as an R package. creDB is designed to enable users to easily consider many different genome-wide sets of enhancers in their bioinformatics analyses and to evaluate the robustness of their findings. The current version of creDB contains more than 3.5 million enhancers and is available online (see Web Resources).

## DISCUSSION

Accurate enhancer identification is a challenging problem, and recent efforts have produced a variety of experimental and computational approaches. Each method, either explicitly or implicitly, represents a different perspective on what constitutes an enhancer and which genomic features are most informative about enhancer activity. The lack of comprehensive genome-wide “gold standard” enhancer sets makes comparisons and evaluation challenging. Thus, we compared existing strategies with respect to one another and to proxies for regulatory function. All pairs of enhancer sets overlap more than expected by chance, but we found substantial differences in the genomic, evolutionary, and functional characteristics of identified enhancers *within* similar tissues and cell types. Enhancer sets vary significantly in their overlap with conserved genomic elements, GWAS loci, and eQTL. Furthermore, the majority of GWAS loci and eQTL have inconsistent evidence of enhancer function across enhancer sets. In addition, regions identified as enhancers by multiple methods *do not* have significantly stronger evidence of regulatory function.

Because enhancer identification strategies have such substantial differences, one strategy cannot and should not be used as a proxy for another. Using different strategies can yield substantially different biological interpretations and conclusions, e.g., about the gene regulatory potential of a SNP or the degree of evolutionary constraint on enhancers. This is particularly important, given that studies of gene regulation commonly use only a single approach to identify enhancers. For example, GWAS have identified thousands of non-coding loci associated with risk for complex disease, and a common first step in the interpretation of a trait-associated locus is to view it in the context of genome-wide maps of regulatory enhancer function^61–63,65,68,73–75^. Thus, our findings complicate the use of annotated enhancers to study the mechanisms of gene regulation and to elucidate the molecular underpinnings of disease, most notably in non-coding variant prioritization^32,76,77^.

Our main goal was to evaluate the congruence of the diverse strategies in use today. Given their differences in assumptions, motivations and protocols, it is not surprising that different assays and algorithms identify somewhat different sets of enhancers; however, the degree of difference we observed is striking. Individual genetic variation may explain some of this discordance. Previous work shows that chromatin states associated with weak enhancer activity exhibit some variation between individuals, and QTL associated with changes in epigenetic modifications leading to variation in enhancer activity between individuals have been identified^78,79^. However, the proportion of epigenetic modifications that are variable across individuals is estimated to be small (1–15%)^80^, and thus is unlikely to be the main cause of the lack of agreement we observe between methods, in particular for enhancer sets defined from cell lines.

The consistent lack of agreement between methods demonstrates that many working definitions of “enhancer” have low overlap. Focusing on functional annotations, we find agreement between methods about basic functions, but substantial differences in more specific annotations. This suggests that different strategies contribute unique information towards the identification of functionally important enhancers. Our results argue that, given the lack of a clear gold standard and the substantial disagreement between strategies, it does not make sense to identify a single “best” method given current knowledge. In general, enhancers defined by FANTOM have modestly more enrichment for proxies of functional activity than other methods, but this comes at the expense of low sensitivity (e.g., Figures S6B, S7B).

In light of this complexity, what should we do? First, we must resist the convenience of ignoring it. When interpreting non-coding variants of interest or characterizing the enhancer landscape in a new biological context, we must be mindful that using a single identification strategy is insufficient to comprehensively catalog enhancers. Different assays and algorithms have different attributes, and we suggest employing a range of approaches to obtain a robust view of the regulatory landscape. To facilitate this, we have developed creDB, a comprehensive, easily queried database of over 3.5 million diverse enhancer annotations. However, simply focusing on variants with multiple lines of evidence of enhancer activity will not solve the problem. We need more sophisticated statistical models of enhancers and their properties in order to interpret non-coding variants of interest. Previous work has shown that integrating diverse genomic, evolutionary, and functional data can improve the ability to distinguish validated enhancers from the genomic background^31^, but obtaining a concordant and functionally relevant set of enhancers remains challenging. We are hopeful that new experimental techniques and biologically motivated machine learning methods for integrating different definitions of enhancers will yield more consistent and specific annotations of regions with regulatory functions.

Second, our study highlights the need for more refined models of the architecture and dynamics of *cis*-regulatory regions. Many different classes of regions with enhancer-like regulatory activities have been discovered^4,14,20,32,39,81,82^. We argue that collapsing the diversity of vertebrate distal gene regulatory regions into a single category is overly restrictive. Simply calling all of the regions identified by these diverse approaches “enhancers” obscures functionally relevant complexity and creates false dichotomies. While there is some appreciation of this subtlety, there is still a need for more precise terminology and improved statistical and functional models of the diversity of cis-regulatory “enhancer-like” sequences. Given this diversity, we should not expect all results to be robust to the enhancer identification strategy used.

Finally, we believe that ignoring enhancer diversity impedes research progress and replication, since “what we talk about when we talk about enhancers” includes diverse sequence elements across an incompletely understood spectrum, all of which are likely important for proper gene expression. Efforts to stratify enhancers into different classes, such as poised and latent, are steps in the right direction, but are likely too coarse given our incomplete current knowledge. We suspect that a more flexible model of distal regulatory regions is appropriate, with some displaying promoter-like sequence architectures and modifications and others with distinct regulatory properties in multiple, potentially uncharacterized, dimensions^2,83,84^. Consistent and specific definitions of the spectrum of regulatory activity and architecture are necessary for further progress in enhancer identification, successful replication, and accurate genome annotation. In the interim, we must remember that genome-wide enhancer sets generated by current approaches should be treated as what they are—incomplete snapshots of a dynamic process.

## MATERIAL AND METHODS

In this section, we first describe the different enhancer identification strategies that we consider. We then describe how we obtained various annotations of functionally relevant attributes for these enhancers. Finally, we describe the analytical approaches used to compare the enhancer sets to one another in terms of their genomic locations and annotations.

### Enhancer identification methods

Here, we summarize how we defined human enhancer sets across four biological contexts. All analyses were performed in the context of the GRCh37/hg19 build of the human genome. We used TSS definitions from Ensembl v75 (GRCh37.p13).

We downloaded broad peak ChIP-seq data for three histone modifications, H3K27ac, H3K4me1, and H3K4me3 from the ENCODE project^33^ for two cell lines, K562 and Gm12878, and from the Roadmap Epigenomics Consortium^34^ for two primary tissues, liver and heart. Also from the ENCODE Encyclopedia (version 3.0), we downloaded the “enhancer-like” annotations; these combine DHSs and H3K27ac ChIP-seq peaks using an unsupervised machine learning model. We retrieved ChromHMM enhancer predictions^35^ for the K562 and Gm12878 cell lines from the models trained on ENCODE data^15^. We downloaded ChromHMM predictions for liver and heart tissues from the 15-state segmentation performed by the Roadmap Epigenomics Consortium. For all ChromHMM sets, we combined the weak and strong enhancer states. We considered two enhancer sets for K562 and Gm12878 based on supervised machine learning techniques—one described in Yip et al. 2012^36^, and the other in Ho et al. 2014^37^. Yip12 predicted ‘binding active regions’ (BARs) using DNA accessibility and histone modification data; the positive set contained BARs overlapping a ‘transcription-related factor’ (TRF), and the negative set contained BARs with no TRF peaks. The predicted regions were filtered using other genomic characteristics to determine the final set of enhancers^36^. Ho14 used H3K4me1 and H3K4me3 ChIP-seq peaks in conjunction with DHSs and p300 binding sites to predict regions with regulatory activity both distal and proximal to TSSs. The distal regulatory elements make up their published enhancer set^37^. We downloaded enhancer regions predicted by the FANTOM consortium for the four sample types analyzed^26^. We also downloaded enhancer predictions in liver from Villar et al. 2015^38^.

To represent enhancer identification strategies in common use, we created three additional enhancer sets for this study using histone modification ChIP-seq peaks and DNase-seq peaks downloaded from ENCODE and Roadmap Epigenomics. The H3K27acPlusH3K4me1 track is a combination of H3K27ac and H3K4me1 ChIP-seq peak files^4,20,39^. If both types of peaks were present (i.e., the regions overlap by at least 50% of the length of one of the regions) the intersection was classified as an enhancer. Similarly, to create the H3K27acMinusH3K4me3 set for each context, we intersected H3K27ac and H3K4me3 ChIP-seq peak files and kept regions where H3K27ac regions did not overlap a H3K4me3 peak by at least 50% of their length. We derived the combination of H3K27ac and H3K4me3 and the 50% overlap criterion from previous studies^14,38,39^. The DNasePlusHistone track is based on the pipeline described in Hay et al. 2016^40^. It combines H3K4me1, H3K4me3, DNaseI hypersensitive sites (DHSs), and transcription start site (TSS) locations. We filtered a set of DHSs, as defined by DNase-seq, for regions with an H3K4me3 / H3K4me1 ratio less than 1, removed regions within 250 bp of a TSS, and called the remaining regions enhancers.

For all enhancer sets, we excluded elements overlapping ENCODE blacklist regions and gaps in the genome assembly^41^. Additionally, due to the presence of extremely long regions in some enhancer sets, likely caused by technical artifacts, we removed any regions more than three standard deviations above or below the mean length of the dataset. This filtering process removed relatively few annotations (Table S10).

### Enhancer attribute data

We downloaded evolutionarily conserved regions defined by PhastCons, a two-state hidden Markov model that defines conserved elements from multiple sequence alignments^42^. We concatenated primate and vertebrate PhastCons elements defined over the UCSC alignment of 45 vertebrates with humans into a single set of conserved genomic regions. We downloaded the full list of 20,458 unique GWAS SNPs from the GWAS Catalog (v1.0, downloaded 08-10-2016)^43^. We also manually curated the set of GWAS SNPs into subsets associated with phenotypes relevant to liver or heart for context-specific analyses (Table S5). We downloaded all GTEx eQTL from the GTEx Portal (v6p, downloaded 09-07-2016)^44^. We concatenated the data from all 44 represented tissues and ran the enrichment analysis on unique eQTL, filtering at four increasingly strict significance thresholds: 10^-6^, 10^-10^, 10^-20^, and 10^-35^. We present the results from the p-value threshold of 10^-10^, although the choice of threshold did not qualitatively alter the results. We also performed separate context-specific analyses on liver and heart specific eQTL from GTEx (p < 10^-10^). To identify other variants tagged by the GWAS SNPs and eQTL, we expanded each set to include SNPs in high LD (r^2^ > 0.9) in individuals of European ancestry from the 1000 Genomes Project (phase 3)^45^.

Experimentally validated enhancer sequences with activity in the heart and all negative enhancer sequences were downloaded from the VISTA enhancer browser (downloaded 11-16-2017)^46^. We also downloaded sequences and Sharpr-MPRA activity levels for 15,720 putative enhancer regions tested for regulatory activity in K562 cells using a massively parallel reporter assay (MPRA)^47^. The Sharpr-MPRA algorithm infers a regulatory score for each base pair in a region using a probabilistic model, with positive scores indicating activating regulatory regions and negative scores indicating repressive regions. Following Ernst et al., we summarized the overall regulatory activity of a given enhancer region as the activity value with the maximum absolute value and classified the enhancer regions into activating (n = 5,373) and repressive (n = 10,347) based on the score’s sign^47^.

### Genomic region overlap and similarity

To quantify genomic similarity, we calculated the base pair overlap between two sets of genomic regions, *A* and *B*, by dividing the number of overlapping base pairs in *A* and *B* by the total number of base pairs in *B*. We also performed this calculation on element-wise level, by counting the number of genomic regions in *B* overlapping regions in *A* by at least 1 bp, and dividing by the number of genomic regions in *B*. We performed both calculations for each pairwise combination of enhancer sets. All overlaps were computed using programs from the BEDtools v2.23.0 suite^48^.

We also evaluated the similarity between pairs of genomic region sets using the Jaccard similarity index. The Jaccard index is defined as the cardinality of the intersection of two sets divided by cardinality of the union. In our analyses, we calculated the Jaccard index at the base pair level. We also computed the relative Jaccard similarity as the observed Jaccard similarity divided by the maximum possible Jaccard similarity for the given sets of genomic regions, i.e., the number of bases in the smaller set divided by the number of bases in the union of the two sets. To visualize overlaps, we plotted heatmaps for pairs of methods using ggplot2 in R^49^.

### Genomic region overlap enrichment analysis

To evaluate whether the observed base pair overlap between pairs of enhancer sets is significantly greater than would be expected by chance, we used a permutation-based approach. We calculated an empirical *p*-value for an observed amount of overlap based on the distribution of overlaps expected under a null model of random placement of length-matched regions throughout the genome. We used the following protocol: let *A* and *B* denote two sets of genomic regions; count the number of bp in *A* that overlap *B*; generate 1,000 random sets of regions that maintain the length distribution of *B*, excluding ENCODE blacklist regions and assembly gaps; count the number of bp in *A* that overlap regions in each of the random sets; compare the observed bp overlap count with the overlap counts from each iteration of the simulation and compute a two-sided empirical p-value. We used the same framework to evaluate element-wise comparisons by counting the number of regions in *A* that overlap *B* rather than the bps. This approach was performed using custom Python scripts and the Genomic Association Tester (GAT)^50^. We note that this measure of overlap significance is not symmetric, and accordingly we confirmed results of our element-wise results for both orderings of the pairs of enhancer sets.

### Enhancer conservation, GWAS catalog SNP, and GTEx eQTL enrichment

In addition to comparing the overlap between pairs of enhancer sets, we also computed enrichment for overlap of evolutionarily conserved regions, GWAS SNPs, and GTEx eQTL with each of the enhancer sets. For conserved elements, we proceeded as described above for comparisons between pairs of enhancer sets, but considered the conserved elements as set *A* and the enhancers as set B. For GWAS tag SNPs, we considered each variant as a region in set *A* and the enhancer regions as set B. We used the same approach for testing all variants in LD (r^2^ > 0.9) with GWAS tag SNPs and for testing enrichment for liver- and heart-specific GWAS tag SNP sets. We also tested for enrichment using only the variant with the maximum number of enhancer set overlaps for each GWAS SNP’s LD block. In this analysis, *A* was the set of variants with maximum enhancer set overlap for each LD block and *B* was the set of enhancers. Enrichments were computed for the eQTL SNP sets using the same strategy as described for GWAS SNPs.

### Enhancer set Gene Ontology annotation and similarity

We used GREAT to find Gene Ontology (GO) annotations enriched among genes nearby the enhancer sets. GREAT assigns each input region to regulatory domains of genes and uses both a binomial and a hypergeometric test to discover significant associations between regions and associated genes’ GO annotation terms^51^. Due to the large number of reported regions in each enhancer set, we considered significance based only on the binomial test with the Bonferroni multiple testing correction (<0.05). We downloaded up to 1,000 significant terms for each enhancer set from the Molecular Function (MF) and Biological Process (BP) GO ontologies. We calculated the similarity between lists of GO terms using the GOSemSim package in R^52^. GOSemSim uses sematic similarity metric that accounts for the hierarchical organization and relatedness of GO terms when calculating the similarity score^53^. For each pair of enhancer sets, we calculated the similarity between their associated GO terms, and converted the resulting similarity matrix into a dissimilarity matrix. We also calculated the number of shared GO terms between pairs of methods and manually compared the top ten significant terms for each enhancer set.

Since enhancers often target genes over long distances, we also considered target predictions generated by the JEME algorithm to assign enhancers to potential target genes in each context^54^. JEME is a two-step process that considers the superset of all enhancers across contexts as well as context-specific biomarkers to make its predictions. By intersecting each enhancer set with the corresponding enhancer-target maps from JEME, we created a set of putatively regulated genes for each method in a given context. We performed GO enrichment analyses on the gene sets using the online tool WebGestalt^55^. We downloaded the top 1,000 significant terms (p < 0.05 after Bonferroni correction) for each enhancer set from the BP and MF GO ontologies, and calculated the pairwise similarity between lists of GO terms using the same semantic similarity metric as above.

### Genomic and functional clustering of enhancer sets

To identify groups of similar enhancers in genomic and functional space, we performed hierarchical clustering on the enhancer sets. For genomic similarity, we converted the pairwise Jaccard similarity to a dissimilarity score by subtracting it from 1 and then clustered the enhancer sets based on these values. For functional similarity, we clustered the lists of GO terms returned by GREAT for each enhancer set. We calculated similarity using the GoSemSim package in R and converted it to dissimilarity by subtracting the similarity score from 1. For both, we used agglomerative hierarchical clustering in R function with the default complete linkage method to iteratively combine clusters^56^. We visualized the cluster results as dendrograms using ggplot2 and dendextend^49,57^. We performed multidimensional scaling (MDS) on the Jaccard and GO term dissimilarity matrices using default options in R^56^.

### Combinatorial analysis of enhancer sets and enrichment for functional signals

We stratified genomic regions by the number of enhancer identification strategies that annotate them in order to determine whether regions predicted to be enhancers by more methods show greater enrichment for three signals of function— evolutionarily conserved base pairs, GWAS SNPs, or GTEx eQTL—compared to regions with less support. We divided all regions predicted by any enhancer identification method in a given context into bins based on the number of methods that predicted it. Some enhancer regions had varying prediction coverage and were split across multiple bins. While infrequent (<3% of regions), we removed all regions less than 10 bp in length since these are unlikely to function as independent enhancers. For each enhancer support bin, from 1 to the number of prediction methods, we calculated the enrichment for overlap with each functional signal using the permutation framework described above. We considered three different proxy sets: evolutionarily conserved base pairs as defined by PhastCons elements, GWAS SNPs, and GTEx eQTL. In each enrichment analysis, the functional signal regions were set *A* and the enhancer regions with a given level of support were set *B*. We report the average enrichment for each enhancer support bin with the empirical 95% confidence intervals.

For enhancer sets with quantitative enhancer-level scores available, we ranked each enhancer by its score, and then analyzed whether regions that have higher scores are more likely to be predicted by other identification methods. We calculated the rank using the ChIP-seq or CAGE-seq signal scores for a subset of methods (H3K27ac, H3K27acPlusH3K4me1, H3K27acMinusH3K4me3, DNasePlusHistone, FANTOM), and the machine learning derived score for EncodeEnhancerlike regions. Within each set, we sorted the enhancer regions by score and assigned ranks starting at 1 for the top-scoring region. We then partitioned the enhancer regions in each set by the number of other enhancer sets that overlap at least one base pair in that region.

### Enhancer database

We created creDB, a database of cis-regulatory elements. It currently contains all 1,371,867 enhancers analyzed in this study and representative sets from other common contexts. It is implemented as an SQLite database with an R interface and distributed as an R package. Our design facilitates the integration of creDB into a wide range of genome-wide bioinformatics workflows, alleviating the vetting and processing that is necessary with flat file downloads. This sets it apart from other efforts that collect enhancers but focus on providing interfaces for searching a small number of candidates or for “browsing” enhancers on a case-by-case basis. For all enhancers, creDB contains information about genomic location, identification strategy, and tissue or cell type of activity. We envision creDB as a growing community resource and encourage other researchers to contribute.

### Data Availability

All data analyzed in this manuscript are available in the creDB R package and database (see Web Resources).

## SUPPLEMENTAL DATA

Supplemental data include seventeen figures (S1-S17) and ten tables (S1-S10).

## ACKNOWLEDGEMENTS

M.L.B. was supported by the National Institutes of Health [T32LM012412]. D.K. was supported by the National Institutes of Health [R01GM115836] and a Basil O’Connor Starter Scholarship from the March of Dimes. J.A.C. was supported by the National Institutes of Health [1R01GM115836, R35GM127087], a March of Dimes Innovation Catalyst award, and the Burroughs Wellcome Fund.

## AUTHOR CONTRIBUTIONS

D.K. and J.A.C. designed and supervised the study. M.L.B. carried out the analyses and made all figures. M.L.B, D.K., and J.A.C. wrote the manuscript. S.C.T. and D.K. designed and implemented the creDB database and software.

## WEB RESOURCES

creDB, http://www.kostkalab.net/software.html

## REFERENCES

1. Shlyueva, D., Stampfel, G. & Stark, A. Transcriptional enhancers: from properties to genome-wide predictions. Nat. Rev. Genet. 15, 272–286 (2014).

2. Andersson, R., Sandelin, A. & Danko, C. G. A unified architecture of transcriptional regulatory elements. Trends in Genetics 31, 426–433 (2015).

3. Ong, C. & Corces, V. Enhancer function: new insights into the regulation of tissue-specific gene expression. Nat. Rev. Genet. 12, 283–93 (2011).

4. Creyghton, M. P. et al. Histone H3K27ac separates active from poised enhancers and predicts developmental state. Proc. Natl. Acad. Sci. U. S. A. 107, 21931–21936 (2010).

5. Maurano, M. T. et al. Systematic Localization of Common Disease-Associated Variation in Regulatory DNA. Science 337, 1190–1195 (2012).

6. Corradin, O. & Scacheri, P. C. Enhancer variants: evaluating functions in common disease. Genome Med. 6, 85 (2014).

7. Sholtis, S. J. & Noonan, J. P. Gene regulation and the origins of human biological uniqueness. Trends Genet. 26, 110–118 (2010).

8. Reilly, S. K. & Noonan, J. P. Evolution of Gene Regulation in Humans. Annu. Rev. Genom. Hum. Genet 1–23 (2016). doi:10.1146/annurev-genom-090314-045935

9. Mack, K. L. & Nachman, M. W. Gene Regulation and Speciation. Trends Genet. 33, 68–80 (2016).

10. Pennacchio, L. A., Bickmore, W., Dean, A., Nobrega, M. A. & Bejerano, G. Enhancers: five essential questions. Nat. Rev. Genet. 14, 288–95 (2013).

11. Kleftogiannis, D., Kalnis, P. & Bajic, V. B. Progress and challenges in bioinformatics approaches for enhancer identification. Brief. Bioinform. 17, 967–979 (2016).

12. Inoue, F. et al. A systematic comparison reveals substantial differences in chromosomal versus episomal encoding of enhancer activity. Genome Res. 27, 38–52 (2017).

13. Inoue, F. & Ahituv, N. Decoding enhancers using massively parallel reporter assays. Genomics 106, 159–164 (2015).

14. Heintzman, N. D. et al. Distinct and predictive chromatin signatures of transcriptional promoters and enhancers in the human genome. Nat. Genet. 39, 311–8 (2007).

15. Hoffman, M. M. et al. Integrative annotation of chromatin elements from ENCODE data. Nucleic Acids Res. 41, 827–841 (2013).

16. Dogan, N. et al. Occupancy by key transcription factors is a more accurate predictor of enhancer activity than histone modifications or chromatin accessibility. Epigenetics Chromatin 8, 1–21 (2015).

17. Inoue, F. et al. A systematic comparison reveals substantial differences in chromosomal versus episomal encoding of enhancer activity. bioRxiv 61606 (2016). doi:10.1101/061606

18. Crawford, G. E. et al. Genome-wide mapping of DNase hypersensitive sites using massively parallel signature sequencing (MPSS). Genome Res. 16, 123–131 (2006).

19. Thurman, R. E. et al. The accessible chromatin landscape of the human genome. Nature 489, 75–82 (2012).

20. Heintzman, N. D. et al. Histone modifications at human enhancers reflect global cell-type-specific gene expression. Nature 459, 108–112 (2009).

21. Woolfe, A. et al. Highly conserved non-coding sequences are associated with vertebrate development. PLoS Biol. 3, (2005).

22. Pennacchio, L. a et al. In vivo enhancer analysis of human conserved non-coding sequences. Nature 444, 499–502 (2006).

23. Visel, A. et al. Ultraconservation identifies a small subset of extremely constrained developmental enhancers. Nat Genet 40, 158–160 (2008).

24. Visel, A. et al. ChIP-seq accurately predicts tissue-specific activity of enhancers. Nature 457, 854–8 (2009).

25. Blow, M. J. et al. ChIP-Seq identification of weakly conserved heart enhancers. Nat. Genet. 42, 806–810 (2010).

26. Andersson, R. et al. An atlas of active enhancers across human cell types and tissues. Nature 507, 455–61 (2014).

27. Li, W., Notani, D. & Rosenfeld, M. G. Enhancers as non-coding RNA transcription units: recent insights and future perspectives. Nat. Rev. Genet. 17, 207–223 (2016).

28. Young, R. S., Kumar, Y., Bickmore, W. A. & Taylor, M. S. Bidirectional transcription marks accessible chromatin and is not specific to enhancers. Genome Biol. (2017). doi:10.1186/s13059-017-1379-8

29. Arner, E. et al. Transcribed enhancers lead waves of coordinated transcription in transitioning mammalian cells. Science 347, 1010 LP-1014 (2015).

30. Taudt, A., Colomé-Tatché, M. & Johannes, F. Genetic sources of population epigenomic variation. Nature Reviews Genetics 17, 319–332 (2016).

31. Erwin, G. D. et al. Integrating diverse datasets improves developmental enhancer prediction. PLoS Comput. Biol. 10, e1003677 (2014).

32. Ernst, J. et al. Mapping and analysis of chromatin state dynamics in nine human cell types. Nature 473, 43–9 (2011).

33. The ENCODE Project Consortium. An Integrated Encyclopedia of DNA Elements in the Human Genome. Nature 489, 57–74 (2012).

34. Roadmap Epigenomics Consortium et al. Integrative analysis of 111 reference human epigenomes. Nature 518, 317–330 (2015).

35. Ernst, J. & Kellis, M. ChromHMM: automating chromatin-state discovery and characterization. Nature Methods 9, 215–6 (2012).

36. Yip, K. Y. K. K. Y. et al. Classification of human genomic regions based on experimentally determined binding sites of more than 100 transcription-related factors. Genome Biol. 13, R48 (2012).

37. Ho, J. W. K. et al. Comparative analysis of metazoan chromatin organization. Nature 512, 449–52 (2014).

38. Villar, D. et al. Enhancer evolution across 20 mammalian species. Cell 160, 554–566 (2015).

39. Rada-Iglesias, A. et al. A unique chromatin signature uncovers early developmental enhancers in humans. Nature 470, 279–83 (2011).

40. Hay, D. et al. Genetic dissection of the α-globin super-enhancer in vivo. Nat. Genet. 1–12 (2016).

41. Kundaje, A. A comprehensive collection of signal artifact blacklist regions in the human genome. … Site/Anshulkundaje/Projects/Blacklists (Last Accessed 30 … (2013).

42. Siepel, A. et al. Evolutionarily conserved elements in vertebrate, insect, worm, and yeast genomes. Genome Res. 15, 1034–1050 (2005).

43. Welter, D. et al. The NHGRI GWAS Catalog, a curated resource of SNP-trait associations. Nucleic Acids Res. 42, (2014).

44. The GTEx Consortium. The Genotype-Tissue Expression (GTEx) project. Nat. Genet. 45, 580–5 (2013).

45. 1000 Genomes Project Consortium et al. A global reference for human genetic variation. Nature 526, 68–74 (2015).

46. Visel, A., Minovitsky, S., Dubchak, I. & Pennacchio, L. A. VISTA Enhancer Browser - A database of tissue-specific human enhancers. Nucleic Acids Res. 35, (2007).

47. Ernst, J. et al. Genome-scale high-resolution mapping of activating and repressive nucleotides in regulatory regions. Nat. Biotechnol. 34, 1180–1190 (2016).

48. Quinlan, A. R. & Hall, I. M. BEDTools: A flexible suite of utilities for comparing genomic features. Bioinformatics 26, 841–842 (2010).

49. Wickham, H. ggplot2. Elegant Graphics for Data Analysis (2009). doi:10.1007/978-0-387-98141-3

50. Heger, A., Webber, C., Goodson, M., Ponting, C. P. & Lunter, G. GAT: A simulation framework for testing the association of genomic intervals. Bioinformatics 29, 2046–2048 (2013).

51. McLean, C. Y. et al. GREAT improves functional interpretation of cis-regulatory regions. Nat. Biotechnol. 28, 495–501 (2010).

52. Yu, G. et al. GOSemSim: An R package for measuring semantic similarity among GO terms and gene products. Bioinformatics 26, 976–978 (2010).

53. Wang, J. Z., Du, Z., Payattakool, R., Yu, P. S. & Chen, C.-F. A new method to measure the semantic similarity of GO terms. Bioinformatics 23, 1274–81 (2007).

54. Cao, Q. et al. Reconstruction of enhancer-target networks in 935 samples of human primary cells, tissues and cell lines. Nat. Genet. 49, 1428–1436 (2017).

55. Wang, J., Vasaikar, S., Shi, Z., Greer, M. & Zhang, B. WebGestalt 2017: a more comprehensive, powerful, flexible and interactive gene set enrichment analysis toolkit. Nucleic Acids Res. 45, W130–W137 (2017).

56. R Core Team. R Core Team (2015). R: A language and environment for statistical computing. R Found. Stat. Comput. Vienna, Austria. URL http://www.R-project.org/. R Foundation for Statistical Computing (2015).

57. Galili, T. dendextend: An R package for visualizing, adjusting and comparing trees of hierarchical clustering. Bioinformatics 31, 3718–3720 (2015).

58. Cotney, J. et al. Chromatin state signatures associated with tissue-specific gene expression and enhancer activity in the embryonic limb. Genome Res. 22, 1069–1080 (2012).

59. Hnisz, D. et al. Super-enhancers in the control of cell identity and disease. Cell 155, 934–47 (2013).

60. Dickel, D. E. et al. Genome-wide compendium and functional assessment of in vivo heart enhancers. Nat. Commun. 7, 12923 (2016).

61. Yao, L., Tak, Y. G., Berman, B. P. & Farnham, P. J. Functional annotation of colon cancer risk SNPs. Nat. Commun. 5, 5114 (2014).

62. Farh, K. K.-H. et al. Genetic and epigenetic fine mapping of causal autoimmune disease variants. Nature 518, 337–43 (2015).

63. Harismendy, O. et al. 9p21 DNA variants associated with coronary artery disease impair interferon-gamma signalling response. Nature 470, 264–268 (2011).

64. Zhernakova, D. V et al. Identification of context-dependent expression quantitative trait loci in whole blood. Nat. Genet. 49, 139–145 (2017).

65. Weinhold, N., Jacobsen, A., Schultz, N., Sander, C. & Lee, W. Genome-wide analysis of noncoding regulatory mutations in cancer. Nat. Genet. 46, 1160–5 (2014).

66. Parker, S. C. J. et al. Chromatin stretch enhancer states drive cell-specific gene regulation and harbor human disease risk variants. Proc. Natl. Acad. Sci. U. S. A. 110, 17921–17926 (2013).

67. Gusev, A. et al. Partitioning heritability of regulatory and cell-type-specific variants across 11 common diseases. Am. J. Hum. Genet. 95, 535–552 (2014).

68. Onengut-Gumuscu, S. et al. Fine mapping of type 1 diabetes susceptibility loci and evidence for colocalization of causal variants with lymphoid gene enhancers. Nat. Genet. 47, 381–6 (2015).

69. Ward, L. D. & Kellis, M. Interpreting noncoding genetic variation in complex traits and human disease. Nat. Biotechnol. 30, 1095–106 (2012).

70. Edwards, S. L., Beesley, J., French, J. D. & Dunning, M. Beyond GWASs: Illuminating the dark road from association to function. American Journal of Human Genetics 93, 779–797 (2013).

71. Ashoor, H., Kleftogiannis, D., Radovanovic, A. & Bajic, V. B. DENdb: Database of integrated human enhancers. Database 2015, (2015).

72. Gao, T. et al. EnhancerAtlas: a resource for enhancer annotation and analysis in 105 human cell/tissue types. Bioinformatics btw495 (2016). doi:10.1093/bioinformatics/btw495

73. Pasquali, L. et al. Pancreatic islet enhancer clusters enriched in type 2 diabetes risk-associated variants. Nat Genet 46, 136–143 (2014).

74. Hazelett, D. J. et al. Comprehensive Functional Annotation of 77 Prostate Cancer Risk Loci. PLoS Genet. 10, e1004102 (2014).

75. Weedon, M. N. et al. Recessive mutations in a distal PTF1A enhancer cause isolated pancreatic agenesis. Nat Genet 46, 61–64 (2014).

76. Tak, Y. G. & Farnham, P. J. Making sense of GWAS: using epigenomics and genome engineering to understand the functional relevance of SNPs in non-coding regions of the human genome. Epigenetics Chromatin 8, 57 (2015).

77. Chatterjee, S. & Ahituv, N. Gene Regulatory Elements, Major Drivers of Human Disease. 1–19 (2017). doi:10.1146/annurev-genom-091416-035537

78. McVicker, G. et al. Identification of genetic variants that affect histone modifications in human cells. Science 342, 747–749 (2013).

79. Kasowski, M. et al. Extensive variation in chromatin states across humans. Science 342, 750–752 (2013).

80. Taudt, A., Colomé-Tatché, M. & Johannes, F. Genetic sources of population epigenomic variation. Nat. Rev. Genet. 17, 319–332 (2016).

81. Pradeepa, M. M. et al. Histone H3 globular domain acetylation identifies a new class of enhancers. Nat. Genet. 48, 681–686 (2016).

82. Zhou, J. & Troyanskaya, O. G. Probabilistic modelling of chromatin code landscape reveals functional diversity of enhancer-like chromatin states. Nat Commun 7, 1–9 (2016).

83. Kim, T. K. & Shiekhattar, R. Architectural and Functional Commonalities between Enhancers and Promoters. Cell 162, 948–959 (2015).

84. Andersson, R. Promoter or enhancer, what’s the difference? Deconstruction of established distinctions and presentation of a unifying model. BioEssays 37, 314–323 (2015).

